# Phosphorylation and ubiquitination independent endocytosis of BRI1

**DOI:** 10.1101/2022.07.11.499622

**Authors:** Lucas Alves Neubus Claus, Derui Liu, Ulrich Hohmann, Nemanja Vukašinović, Roman Pleskot, Jing Liu, Alexei Schiffner, Yvon Jaillais, Guang Wu, Sebastian Wolf, Daniël Van Damme, Michael Hothorn, Eugenia Russinova

**Affiliations:** Department of Plant Biotechnology and Bioinformatics, Ghent University, 9052 Ghent, Belgium; Center for Plant Systems Biology, VIB, 9052 Ghent, Belgium; Structural Plant Biology Laboratory, Department of Botany and Plant Biology, University of Geneva, 1211 Geneva, Switzerland; College of Life Sciences, Shaanxi Normal University, Xi’an, Shaanxi, 710062, P.R. China; Center for Plant Molecular Biology (ZMBP), University of Tübingen, 72076 Tübingen, Germany; Laboratoire Reproduction et Développement des Plantes (RDP), Université de Lyon, Ecole Normale Supérieure de Lyon, Centre National de la Recherche Scientifique (CNRS), Institut National de Recherche pour l’Agriculture, l’Alimentation et l’Environnement (INRAE), 69342 Lyon, France; College of Life Sciences, Shandong Agricultural University, Tai’an 271018, China; Institute of Molecular Biotechnology of the Austrian Academy of Sciences (IMBA) & Institute of Molecular Pathology (IMP), Vienna BioCenter (VBC), 1030 Vienna, Austria; Institute of Experimental Botany, Academy of Sciences of the Czech Republic, 16502 Prague 6, Czech Republic

**Keywords:** BRI1, brassinosteroids, ligand-binding, endocytosis, non-canonical

## Abstract

The brassinosteroid (BR) hormone and its plasma membrane receptor BR INSENSITIVE1 (BRI1) is one of the best-studied receptor-ligand pairs for understanding the interplay between receptor endocytosis and signaling in plants. BR signaling is mainly determined by the plasma membrane pool of BRI1, whereas BRI1 endocytosis ensures signal attenuation. Since BRs are ubiquitously distributed in the plant, the tools available to study BRI1 function without interference from endogenous BRs are limited. Here, we designed a BR-binding-deficient mutant based on protein sequence-structure analysis and homology modeling of BRI1 and its close homologues. This new tool allowed us to re-examine the BRI1 endocytosis and signal attenuation model. We show that despite decreased phosphorylation and ubiquitination, the BR-binding-deficient BRI1 was internalized similar to the wild type form. These results reinforce the hypothesis that BRI1 is internalized via parallel endocytic routes and machineries. In addition, BR-binding-deficient mutant provides opportunities to study non-canonical ligand-independent BRI1 functions.

## Introduction

Brassinosteroids (BRs) are low abundant and ubiquitously distributed plant steroidal hormones that play essential role in growth, development, immunity and responses to stress (Nolan et al., 2020). BR biosynthetic or signaling mutants display severe phenotypes including dwarfism, dark-green leaves, photomorphogenesis in the dark, and late flowering (Nolan et al., 2020). BRs are perceived at the cell surface by a leucine-rich repeat (LRR) receptor kinase BR INSENSITIVE1 (BRI1) (He et al., 2000; Hothorn et al., 2011; Kinoshita et al., 2005; She et al., 2011; Wang et al., 2001). BR binding triggers the dissociation of the inhibitory proteins BRI1 KINASE INHIBITOR1 (BKI1) and BAK1-INTERACTING RECEPTOR-LIKE KINASES3 (BIR3), allowing interaction between BRI1 and its coreceptor BRI1-ASSOCIATED KINASE1 (BAK1), which is required for downstream signaling (Hohmann et al., 2018a; Li et al., 2002; Wang and Chory, 2006). BR signal is conveyed from the cell surface to the nucleus through a sequence of phosphorylation/de-phosphorylation events that activate the transcription factors of the BRASSINAZOLE-RESISTANT1 (BZR1) and BRI1-EMS-SUPPRESSOR1 (BES1)/BZR2 family (Wang et al., 2002; Yin et al., 2002; Chen et al., 2019). BRI1 receptor functions go beyond its canonical BR signaling function as together with the RECEPTOR-LIKE PROTEIN44 (RLP44) and BAK1, BRI1 controls xylem cell fate independently of BRs (Holzwart et al., 2018, 2020).

An important regulatory step in BR signaling is the control of the plasma membrane (PM) pool of BRI1, which is determined by BRI1 endocytosis, recycling, and secretion (Irani et al., 2012; Luo et al., 2015). As a consequence, impaired endocytosis or receptor secretion, enhanced and reduced BR signaling, respectively (Irani et al., 2012; Luo et al., 2015). Several studies have focused on BRI1 dynamics upon ligand binding. Given that exogenous BRs did not change BRI1 internalization dynamics, BRI1 endocytosis was described as ligand-independent (Geldner et al., 2007; Luo et al., 2015; Russinova et al., 2004). Subsequent studies on post-translation modifications (PTM) showed that BRI1 undergoes polyubiquitination that is mediated by the plant U-box (PUB) E3 ubiquitin ligases PUB12 and PUB13 and requires BR binding (Zhou et al., 2018). BRI1 ubiquitination is a signal for BRI1 endocytosis and vacuolar sorting, since disruptions in this process by mutations in either the ubiquitination sites or the E3 ligases translate into accumulation of BRI1 in the PM and consequently BR sensitivity increases (Martins et al., 2015; Zhou et al., 2018).

Available tools used to study the dependence of BRI1 trafficking on ligand-binding are limited and rely on the depletion of the endogenous BRs by using BR biosynthetic mutants or the BR biosynthesis inhibitor, brassinazole (BRZ) (Asami et al., 2000). However, the possibility cannot be excluded that treatment with BRZ might not completely deplete bioactive BRs and that the BR biosynthetic mutant might contain biologically active BR precursors. Furthermore, BR biosynthetic mutants display pleiotropic phenotypes that could lead to general changes in membrane trafficking. Therefore, the use of these tools could hamper analysis of ligand-independent BRI1 dynamics.

Here, we report the characterization of a quintuple (Q) BR binding-deficient BRI1 receptor mutant, designated as BRI1^Q^, that was generated using homology and structure analysis of BRI1 and its three homologues BRI1-LIKE1 (BRL1), BRL2 and BRL3 (Caño-Delgado et al., 2004). This new tool reveled that BRI1 endocytosis is largely independent of BRs and despite strongly decreased phosphorylation and ubiquitination the BRI1^Q^ mutant held normal endocytosis rates. These results reinforce the hypothesis that BRI1 is internalized via parallel endocytic routes and machineries. Moreover, we used the BRI1^Q^ mutant to investigate BR-independent BRI1 functions, such as xylem cell differentiation. We showed that BRI1^Q^ can partially complement the xylem cell fate phenotype of the *bri1* null mutant, suggesting that BRI1^Q^ can be used to study BRI1 non-canonical functions.

## Results and Discussion

### BRI1^Q^ cannot bind BRs

The *Arabidopsis thaliana* (Arabidopsis) genome encodes for three BRI1 homologs, designated BRL1, BRL2 and BRL3, of which BRL2 does not bind BRs (Caño-Delgado et al., 2004; Kinoshita et al., 2005). Sequence analyses of BRI1, BRL1, BRL2 and BRL3 ectodomains (Figure 1A) and examination of the crystal structure of the BRI1 ectodomain in a complex with brassinolide (BL), the most active BR (Hothorn et al., 2011; She et al., 2011; Wang et al., 2001) (Figure 1B-1D), revealed five putative residues important for BR binding, three derived from changes in the BRL2 sequence [tyrosine (Y)597, Y599 and methionine (M)657] and two bulky hydrophobic residues located at a 4 Å distance from the BL molecule in BRI1 [Y642 and phenylalanine (F)681] (Figure 1B and 1C). Y597, Y599 and Y642 map to the inner surface of the BRI1 island domain, forming the distal part of the BR-binding pocket, whereas M657 and F681 are located in the LRR core and establish hydrophobic interactions with the aliphatic BL moiety (Figure 1B and 1C). The identified residues were mutated to the corresponding ones in BRL2 either individually (BRI1^Y599F^ and BRI1^M657E^) or in a combination (BRI1^Y597M/Y599F/M657E^) or to alanine (A) (BRI1^Y597A/Y599A/M657A^). Finally, a quintuple BRI1^Y597M/Y599F/Y642A/M657E/F681A^ version was generated and designated as BRI1^Q^. Next, the binding kinetics of BL to the BRI1^Q^ ectodomain was determined by grating-coupled interferometry (GCI) (Figure 2B). As controls, the ectodomains of wild type BRI1 and that of the previously characterized in *bri1-6* mutant that carries the glycine (G) 644 to asparagine (D) missense mutation (BRI1^G644D^) were included (Hohmann et al., 2018a; Hothorn et al., 2011; Kinoshita et al., 2005; Noguchi et al., 1999; Wang et al., 2001). Analytical size-exclusion chromatography and right-angle light scattering experiments confirmed that all BRI1 variants were monodisperse, suggesting that mutations in the BR-binding pocket do not affect the overall shape and oligomeric state of the BRI1 ectodomain (Figure S1). The GCI experiments revealed that BRI1 bound BL with a dissociation constant (K_*D*_) of ∼10 nM as previously reported (Hohmann et al., 2018a; Wang et al., 2001), BRI1^Q^ did not bind BL, whereas the BRI1^G644D^ mutant displayed a strongly reduced BL-binding capacity with a K_*D*_ of ∼11.6 μM.

**Figure 1.**
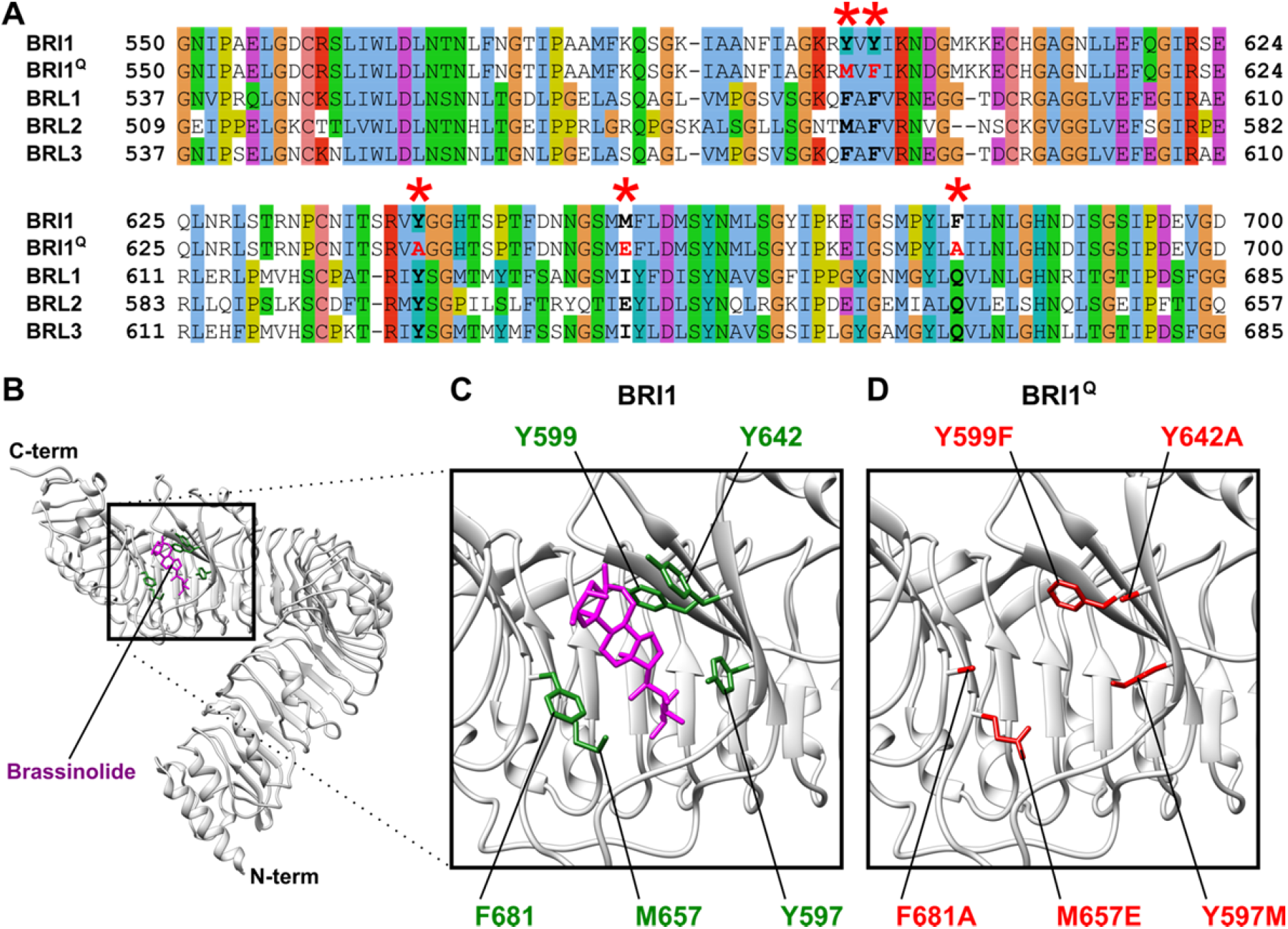
Selection of the essential residues for BR binding in the BRI1 ectodomain. (A) Sequence alignment of the wild type BRI1 ectodomain (only the region corresponding to BRI1 550-700 is shown) with that of BRL1, BRL2 and BRL3 and BRI1^Q^ with the mutated residues highlighted in red. The mutated residues are marked with an asterisk. (B) The BRI1 ectodomain structure in a complex with brassinolide (BL) (Protein Data Bank ID 3RJ0). (C and D) The five residues selected are indicated as single-letter abbreviation with their locations, either colored in green for wild type BRI1 or in red for BRI1^Q^. The figures (B-D) were generated with UCSF Chimera where the BL molecule is shown in a balls-and-sticks representation and colored in purple.

**Figure 2.**
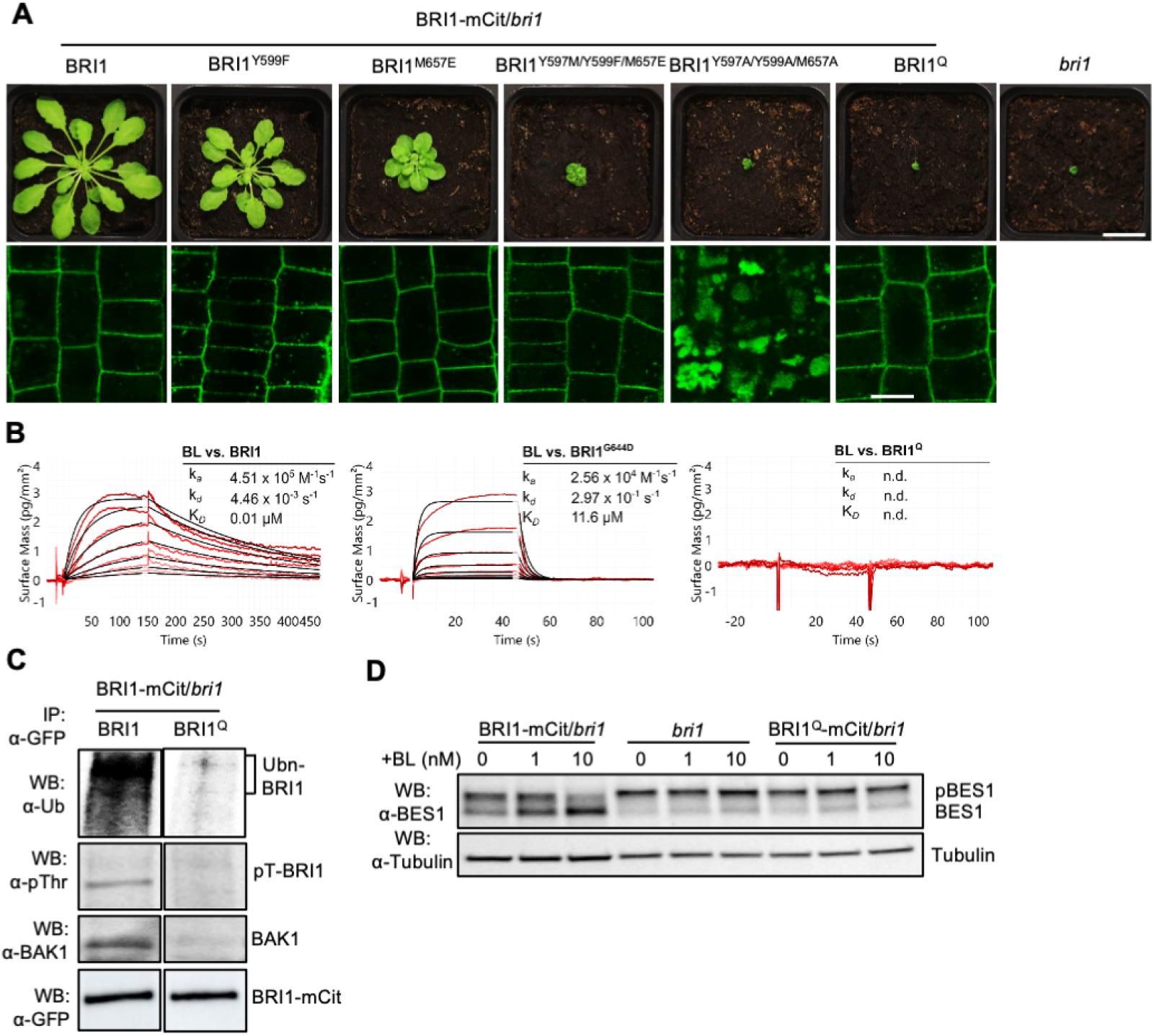
BRI1^Q^ cannot bind BL. (A) Phenotypes (upper panel) and subcellular localizations (lower panel) of homozygous *bri1* mutant transgenic plants expressing the indicated BRI1 isoform mutations grown in short-day cycle for 6 weeks. The quintuple BRI1^Y597M/Y599F/Y642A/M657E/F681A^ mutation is designated BRI1^Q^. Epidermal root meristem cells of 5-day-old seedlings were imaged. Scale bars, 2 cm (upper panel) and 10 μm (lower panel). (B) Binding kinetics for BL vs wild-type BRI1, BRI1^G644D^ and BRI1^Q^ as obtained from grating-coupled interferometry (GCI). Sensograms with recorded data are shown in red with the respective fits in black (when applicable) and include the corresponding association rate constant (k_*a*_), dissociation rate constant (k_*d*_) and dissociation constant (K_*D*_). (C) BRI1-mCit and BRI1^Q^-mCit phosphorylation and ubiquitination state and interaction with BAK1 were tested by isolation of microsomal fractions of 5-day-old seedlings followed by immunoprecipitation (IP) and Western blot (WB) analysis with α-ubiquitin (α-Ub), α-pThreonine (α-pThr) and α-BAK1 antibodies. (D) BES1 phosphorylation state assessed in 5-day-old seedlings treated with DMSO, 1 nM or 10 nM BL for 1 h were subjected to WB analysis with the α-BES1 antibody; α-tubulin was used as loading control.

After confirmation that BRI1^Q^ cannot bind BL, mutated full-length BRI1 versions fused to mCitrine (mCit) were expressed in the *bri1* null mutant from the native promoter and plants with similar protein expression levels of the transgenes were selected (Figures 2A and S2A). BRI1^Y599F^-mCit, BRI1^M657E^-mCit and BRI1^Y597M/Y599F/M657E^-mCit partially complemented the *bri1* dwarf phenotype and localized in the PM and in intracellular punctate structures, similar to the wild type BRI1 (Geldner et al., 2007; Russinova et al., 2004) (Figure 2A). By contrast, BRI1^Y597A/Y599A/M657A^-mCit and BRI1^Q^-mCit did not complement the *bri1* mutant (Figure 2A). The reason for the lack of *bri1* complementation by BRI1^Y597A/Y599A/M657A^-mCit was the aberrant localization of this BRI1 mutant isoform in the vacuole. BRI1^Q^-mCit, however, exhibited the correct BRI1 localization in the PM and in intracellular punctate structures. Hence, the absence of BRI1^Q^ functionality corroborates the *in vitro* ligand-binding deficiency results (Figure 2B). To further characterize the BRI1^Q^-mCit line, we tested the BL-induced BRI1 PTMs. It is well established that after ligand binding, BRI1 heterodimerizes with its co-receptor BAK1 and undergoes PTMs such as phosphorylation and ubiquitination (Belkhadir and Jaillais, 2015; Martins et al., 2015; Zhou et al., 2018). Moreover, BL treatment promotes the dephosphorylation of the transcription factor BES1 in a dose-dependent manner, which is frequently used as a BR signaling indicator (Yin et al., 2002). In agreement with the impaired BL binding, BRI1^Q^-mCit had no detectable phosphorylation, ubiquitination, did not interact with BAK1 and did not promote BES1 dephosphorylation after BL treatment (Figure 2C and 2D). These findings indicate that BRI1^Q^ is unable to perceive BRs.

### Endocytosis of BRI1 is independent of BR binding

The lack of BL binding in the BRI1^Q^ mutant provides a powerful tool to investigate different aspects of BRI1 regulation, including endocytosis, without interference from BRs. Although initially BRI1 endocytosis had been described as ligand independent (Geldner et al., 2007; Russinova et al., 2004), later BR perception has been demonstrated to promote BRI1 ubiquitination (Zhou et al., 2018), which assists BRI1 internalization and vacuolar targeting (Luo et al., 2022; Martins et al., 2015; Zhou et al., 2018). We revisited the BRI1 ligand-dependent endocytosis model using BRI1^Q^ by evaluating the internalization of BRI1^Q^-mCit in root meristem epidermal cells (Figure 3A and 3B). The PM pool of BRI1 is regulated by secretion, recycling, and endocytosis. To avoid interference of the newly synthesized and secreted BRI1, we analyzed 5-day-old plants expressing BRI1^Q^-mCit treated with 50 µM of the protein synthesis inhibitor cycloheximide (CHX) for 1.5 h. Interestingly, the PM *vs*. cytoplasm fluorescence intensity, did not significantly differ between BRI1^Q^-mCit and the control BRI1-mCit, both in *bri1* null background (Figures 3A and 3B, upper panel). In addition to the CHX treatment, we applied Brefeldin A (BFA), an inhibitor of endosomal trafficking that is widely used to visualize endocytosis (Geldner et al., 2003). In Arabidopsis roots, BFA treatment promotes the formation of BFA bodies, composed of large aggregation of trans-Golgi network/early endosome (TGN/EE) compartments (Geldner et al., 2003; Lam et al., 2009). When combining CHX (50 µM, 1.5 h) with BFA (50 µM, 30 min), both BRI1-mCit and BRI1^Q^-mCit accumulated in similar size BFA bodies (Figure 3A and 3B, lower panel). These results were in agreement with the measurements of the PM *vs*. cytoplasm fluorescence intensity in BRI1^Q^-mCit (Figure 3A and 3B, upper panel). Comparable PM *vs*. cytoplasm fluorescence intensity ratios and BFA body size were also obtained when BRI1^Q^-mCit was introduced into the Columbia-0 (Col-0) wild type to avoid artefacts in quantitative microscopy due to the strong dwarfism of the *bri1* mutant (Figure 3C and 3D, S2B).

**Figure 3.**
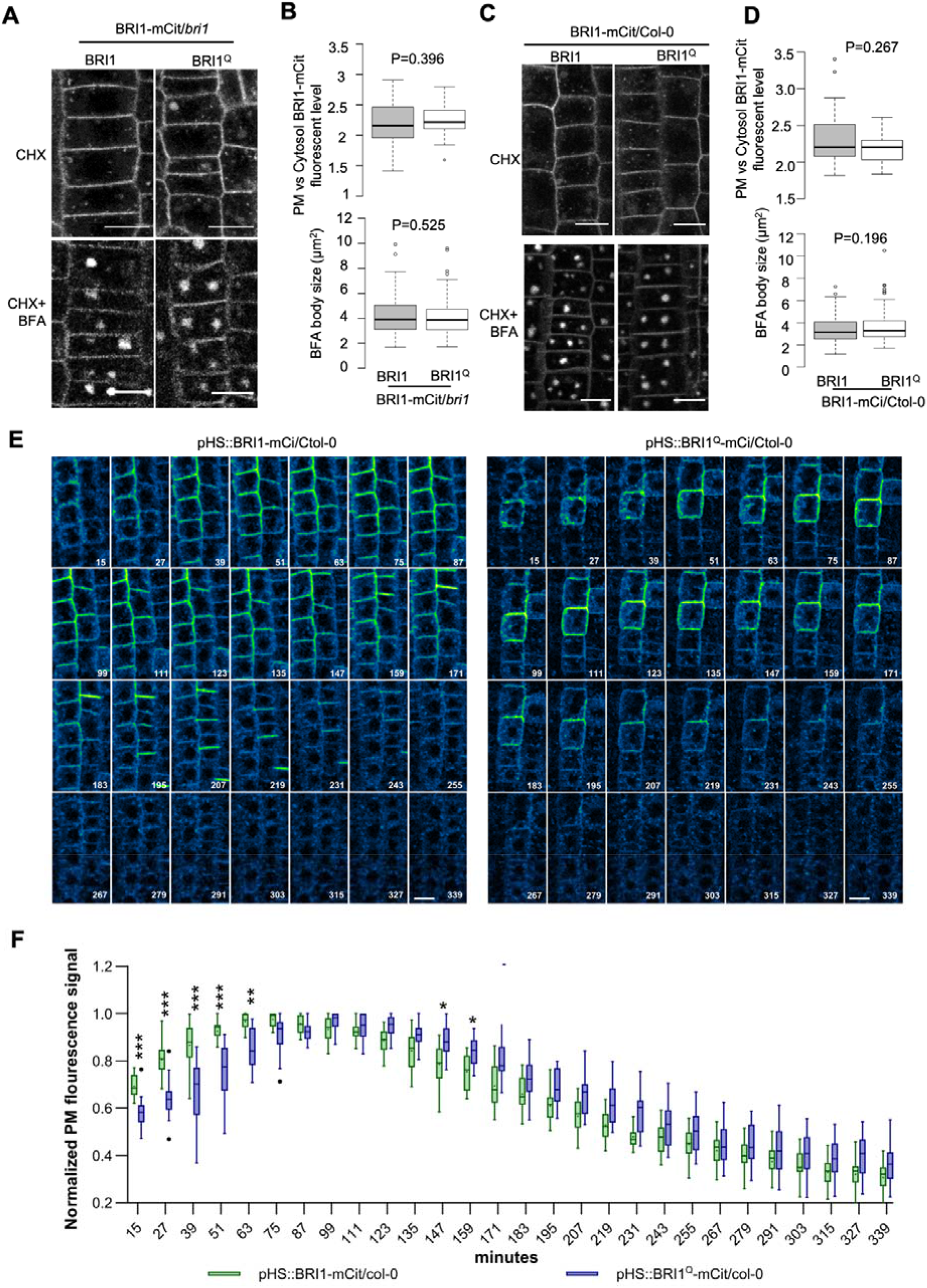
BRI1^Q^ endocytosis is independent of ligand binding. (A and C) Representative confocal images of epidermal root meristem cells of 5-day-old seedlings expressing BRI1-mCit or BRI1^Q^-mCit treated with CHX (50 µM) for 1.5 h or pretreated with CHX for 1 h, followed by treatment for 30 min with CHX and BFA (50 μM). Scale bar, 10 µm. (B and D) Plasma membrane (PM) *vs* cytosol BRI1-mCitrine fluorescence and BFA body size. For each line, 15 cells from at least 5 seedlings were measured. Statistical analysis was performed using Mann Whitney test. (E and F) Time series analysis of meristem epidermal cells of 5-day-old seedling expressing BRI1-mCit or BRI1^Q^-mCit after heat induction (1 h at 37°C). Images were taken with a vertical confocal microscope with a 12 min interval between the frames. Scale bar, 10 µm. Plasma membrane signal intensity of the same cells was quantified for all time points and the signal peak was set as 1. Four roots and 3 to 5 cells were measured per genotype. Asterisks indicate statistically significant differences, * *P*<0.05, ** *P*<0.01, *** *P*<0.001, Two-way ANOVA post hoc Sidak’s multiple comparisons test.

Despite the widespread use of CHX and BFA treatments to analyze endocytosis in plant cells, it cannot be excluded that that the chemical treatments might cause pleiotropic effects (Oksvold et al., 2012; Smith et al., 2014). To circumvent this problem, we expressed BRI1-mCit and BRI1^Q^-mCit under the control of the heat shock-inducible promoter (*pHS*) in the Col-0 background and studied the protein internalization during the recovery phase following induction at 37ºC for 1 h. Firstly, we selected transgenic lines expressing *pHS::BRI1-mCit* and *pHS::BRI1*^*Q*^*-mCit* with similar expression levels following the heat induction (37ºC) for 1 h (Figure S2C). Taking advantage of a vertical confocal microscope setup equipped with the TipTracker software (von Wangenheim et al., 2017) that allows the monitoring of growing root tips over time, we observed that BRI1^Q^-mCit reached the PM a little later than the BRI1-mCit. However, after a signal intensity peak in the PM, between 87 and 99 min, the internalization rate of BRI1-mCit and BRI1^Q^-mCit was very similar. Thus, our findings further confirm previous reports (Geldner et al., 2007; Irani et al., 2012; Russinova et al., 2004) that the BRI1 internalization is largely independent of BR binding. After ligand binding and interaction with BAK1, BRI1 is phosphorylated and ubiquitinated, both essential for receptor internalization (Martins et al., 2015; Zhou et al., 2018). However, although BRI1^Q^ lacked both PTMs, it still had a normal endocytosis. These results reinforce the hypothesis that BRI1 is internalized via more than one endocytic route and probably via different mechanisms. For instance, the BRI1 internalization has been demonstrated to partially depend on both the classical clathrin Adaptor Protein 2 (AP-2) complex that binds to a canonical YXXΦ endocytic motif in BRI1 (Liu et al., 2020) and on ubiquitin recognition machinery, since endocytosis of the ubiquitin-deficient BRI1^25KR^-mCit or BRI1-mCit in the *pub12 pub13* double mutant was not completely abolished (Martins et al., 2015; Zhou et al., 2018). Similar observations were reported for the borate exporter BOR1, in which AP-2-dependent and AP-2-independent endocytic routes had been activated by low and high borate concentration, respectively (Yoshinari et al., 2019). In mammals, the well-studied epidermal growth factor receptor (EGFR) is also internalized via multiple endocytic routes, including canonical ligand-dependent clathrin-mediated endocytosis and clathrin-independent endocytosis route, which both depend on the ligand concentration (Zhou and Sakurai, 2022), and a ligand-independent route where EGFR endocytosis is induced by stress conditions and does not require kinase activity or ubiquitination (Metz et al., 2021). Taken together, our results show that the BRI1 internalization is not abolished in the BR binding-deficient mutant, similarly to BRI1 YXXΦ endocytic motif mutants (Liu et al., 2020), the BRI1 ubiquitination-deficient mutants (Martins et al., 2015; Zhou et al., 2018) and the BRI1 endocytosis in AP-2 mutants(Di Rubbo et al., 2013; Gadeyne et al., 2014). It remains to be established which signals or conditions trigger these different types of BRI1 endocytosis.

### BRI1^Q^ can partially complement the xylem phenotype of *bri1*

In addition to its primary role in perceiving BRs, recent studies suggest that BRI1 might also have non-canonical functions in sensing cell wall integrity. After disturbance of the cell wall integrity by the inhibition of the pectin de-methyl esterification, BRI1 is recruited together with the RLP44 and BAK1 to activate a BR signaling for a compensatory feedback loop to remodel the cell wall (Wolf et al., 2012). Besides cell wall integrity monitoring, RLP44 is also implicated in controlling xylem cell fate through the phytosulfokine (PSK) signaling (Holzwart et al., 2018). Interestingly, BRI1 and BAK1 are also necessary to regulate the vasculature cell fate, but independently of BRs, because BR biosynthesis mutants have no ectopic xylem in the procambial position present in the *rlp44* and *bri1* mutants (Holzwart et al., 2018, 2020). To test whether BRI1^Q^ retains its non-BRs receptor functions, we examined if BRI1^Q^ could still interact with RLP44 (Holzwart et al., 2018). RLP44-RFP was transiently co-expressed with either BRI1-GFP or BRI1^Q^-GFP in *Nicotiana benthamiana* leaves and co-immunoprecipitation (Co-IP) assay reveled that RLP44-RFP was co-purified with both BRI1-GFP and BRI1^Q^-GFP but not with the negative control, indicating that BRI1^Q^, like the wild type BRI1, can form a complex with RLP44 (Figure 4A). Furthermore, confocal analysis of tobacco leaves transiently expressing BRI1-GFP, BRI1^Q^-GFP and RLP44-RFP show that both BRI1 and BRI1^Q^ colocalize with RLP44 in dynamic punctate structures (Figure 4B). The intracellular punctate structures containing RLP44-RFP and BRI1-GFP or BRI1^Q^-GFP are probably endosomes, because BRI1 is a *bona fide* endosomal PM cargo (Geldner et al., 2007; Russinova et al., 2004). Moreover, RLP44 had already been shown to localize in endosomal structures in Arabidopsis roots (Wolf et al., 2014). Finally, we tested whether BRI1^Q^-mCit could recover the BR-independent xylem cell fate phenotype of the *bri1* null mutants (Holzwart et al., 2018, 2020). Indeed, BRI1^Q^-mCit could partially complement the ectopic number of xylem cells present in the *bri1* mutant (Figure 4). Collectively, our result provides evidence that BRI1^Q^ could still be active in BR-independent pathways.

**Figure 4.**
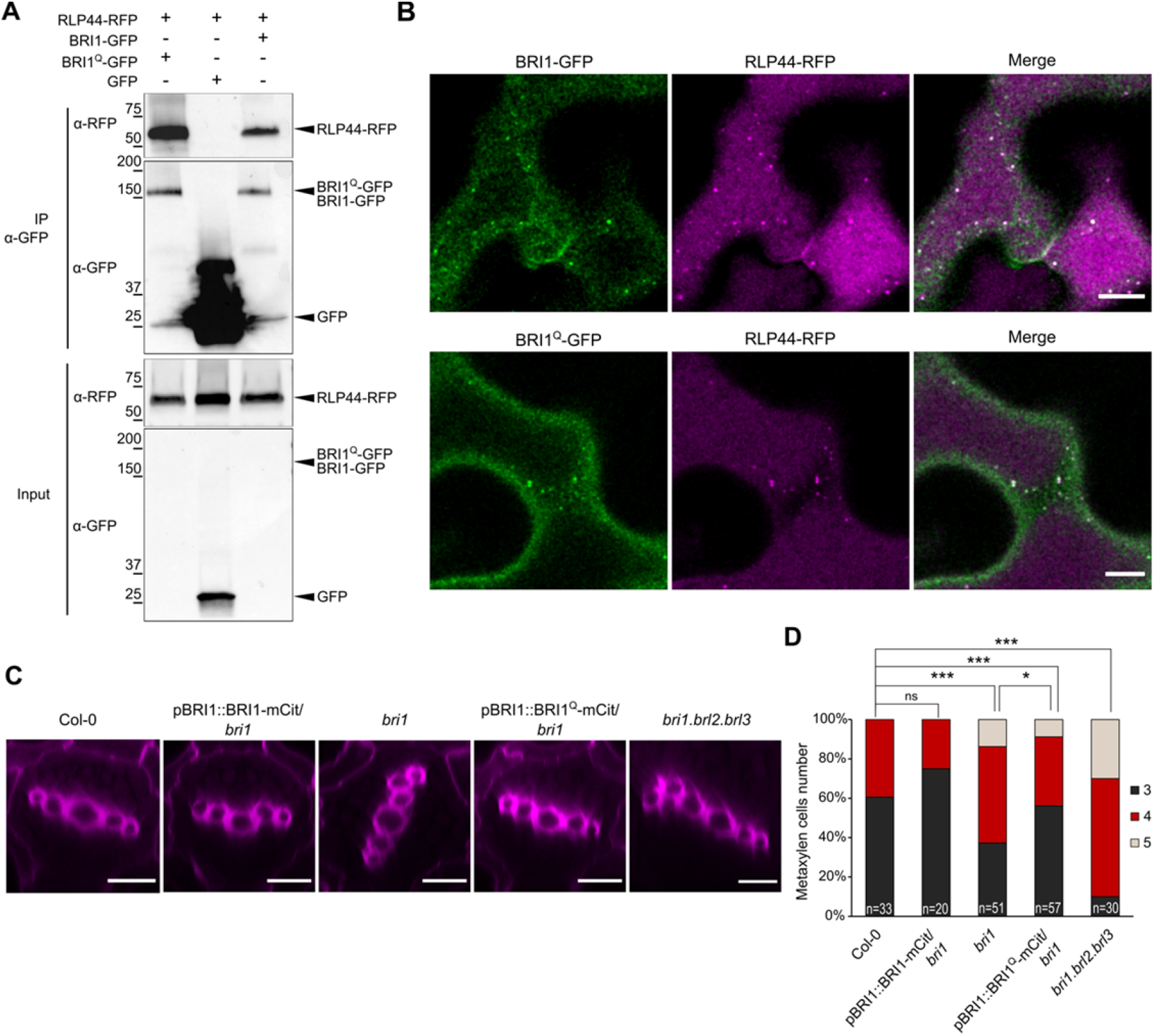
BRI1^Q^ partially recover *bri1* xylem cell fate phenotype. (A) Coimmunoprecipitation of RLP44-RFP transiently co-expressed with BRI1-GFP, BRI1^Q^-GFP, or free GFP (negative control) in *Nicotiana benthamiana* leaf epidermal cells. Proteins were extracted (Input) and immunoprecipitated (IP) by means of magnetic GFP beads. The immunoblot was done with α-GFP and α-RFP antibodies. (B) Co-localization of RLP44-RFP with BRI1-GFP or BRI1^Q^-GFP in subcortical discrete punctate structures in *Nicotiana benthamiana* leaf epidermal cells. Scale bar, 10 µm. (C) 7-day-old roots stained with Basic Fuchsin for the visualization of xylem cells. Scale bar, 10 µm. (D) Frequency quantification of roots with the indicated number of metaxylem cells. Asterisks indicate statistically significant differences, * *P*<0.05, ** *P*<0.01, *** *P*<0.001, Chi-square test. *n*=20-57 as indicated.

In conclusion, analyses of the crystal structure of the ectodomain of BR receptors and homology modeling allowed us to create a BR binding-deficient BRI1 mutant, BRI1^Q^. Characterization of BRI1^Q^ showed that it displays a clear *bri1*-like phenotype and is unable to respond to exogenous BRs. By means of BRI1^Q^ as a tool to study BRI1 endocytosis and in agreement with previous observations (Geldner et al., 2007; Irani et al., 2012; Russinova et al., 2004), we conclude that BRI1 internalization can occur without BR binding. Moreover, the BR-binding-deficient BRI1 mutant might provide opportunities to discover additional non-canonical ligand-independent BRI1 functions in vasculature development and other processes (Graeff et al., 2020; Holzwart et al., 2018, 2020).

## Materials and Methods

### Plant materials, growth conditions and treatments

The experimental model used in this study is Arabidopsis. The wild type used is the *Arabidopsis thaliana* (L.) Heynh. (accession Columbia-0 [Col-0]). For phenotypic analysis, plants were grown in soil in a growth chamber at 22°C, 58% relative humidity, and a 16-h light/8-h dark photoperiod for 6 weeks. The Arabidopsis seeds were surface-sterilized with chlorine gas, and then placed on plates with half-strength Murashige and Skoog medium (½MS) containing 0.5% (w/v) sucrose, 0.8% (w/v) agar, and 2.5 mM methyl ester sulfonate at pH 5.7. After vernalization for 2 days at 4°C, the plates were moved to the growth chamber under a 16-h/8-h light/dark cycle. Wild-type tobacco (*Nicotiana benthamiana*) plants were grown in the greenhouse under a normal 14-h light/10-h dark regime at 25°C. For the microsomal protein preparation, plants were grown for 6 days on plates. For the BRI1 internalization assay and the BRI1 transcript analysis, plants were grown for 5 or 7 days on plates. MG-132 (10 mM stock in dimethylsulfoxide [DMSO]), BFA (50 mM stock in DMSO) and CHX (50 mM stock in DMSO) were used at the concentrations indicated in the figure legends.

### Vector construction and plant transformation

The BRI1-coding region without the stop codon was cloned into *pMD19-T* (simple) (Takara Biotechnology) to generate *pMD19-BRI1* that was used as a template to generate the mutants BRI1^Y599F^, BRI1^M657E^, BRI1^Y597M/Y599F/M657E^, BRI1^Y597A/Y599A/M657A^, and BRI1^Y597M/Y599F/Y642A/M657E/F681A^ by overlapping PCR and subcloned into *pDONR221* to generate *pDONRP1P2-BRI1* (with mutations). The primers used to generate the BRI1 mutations are listed in Table S1. The destination vectors were generated by recombining *pK7m34GW, pB7m34GW, pDONRP4P1r-pBRI1, pDONRP4P1r-pHS* (Marquès-Bueno et al., 2016) *pDONRP4P1r-pBRI1, pDONRP4P1r-pHS, pDONR221-BRI1*, and *pDONRP2rP3-mCit* (Martins et al., 2015). The resulting constructs were transformed into the heterozygous *bri1* null mutant (GABI_134E10) (Jaillais et al., 2011) or into Col-0 plants by floral dip. For transient expression in tobacco, *pDONR221-BRI1* and *pDONR221-BRI1*^*Q*^ were recombined in pK7FWG2 that contained the 35S promoter and C-terminal GFP. The Gateway technology (Invitrogen) was used for cloning.

### Western blot analysis and immunoprecipitation

For the BRI1 expression assay, 5-day-old seedlings were homogenized in liquid nitrogen. Total proteins were extracted with a buffer containing 20 mM Tris-HCl, pH 7.5, 150 mM NaCl, 1% (w/v) sodium dodecyl sulfate (SDS), 100 mM dithiothreitol (DTT), and ethylenediaminetetraacetic acid (EDTA)-free protease inhibitor cocktail cOmplete (Roche). For blocking and antibody dilutions, 3% (w/v) bovine serum albumin (BSA) powder in 0.2% (v/v) Tris-buffered saline-containing Tween-20 was used. For the microsomal fraction isolation, 6-day-old seedlings treated with 50 µM MG-132 for 5 h were ground in liquid nitrogen and resuspended in ice-cold sucrose buffer (100 mM Tris [pH 7.5], 810 mM sucrose, 5% [v/v] glycerol, 10 mM EDTA [pH 8.0], 10 mM ethyleneglycoltetraacetic acid [EGTA, pH 8.0], 5 mM KCl, protease inhibitor [Sigma-Aldrich], and phosphatase inhibitor [Sigma-Aldrich]). The homogenate was transferred to polyvinyl polypyrrolidone (PVPP) pellets, mixed, and left to rest for 5 min. Samples were centrifuged for 5 min at 600*g* at 4°C. The supernatant was collected. The extraction was repeated for two more times. The supernatant was filtered through a Miracloth mesh. The same amount of water was added to the clear supernatant and centrifuged at 21,000*g* for 2 h at 4°C to pellet microsomes (Abas and Luschnig, 2010). The pellet was resuspended in immunoprecipitation buffer (25 mM Tris, pH 7.5, 150 mM NaCl, 0.1% [w/v] SDS, protease inhibitor, and phosphatase inhibitor). Immunoprecipitations were carried out on solubilized microsomal proteins with GFP-Trap-MA (Chromotek) according to the manufacturer’s protocol. For protein detection, the following antibodies were used: monoclonal α-GFP horseradish peroxidase-coupled (1/5,000; Miltenyi Biotech), monoclonal α-tubulin (1/10,000; Sigma-Aldrich), α-ubiquitin (Ub) P4D1 (1/2,500; Millipore), α-pThr (1/2,000; Cell Signaling), α-BES1 (Yin et al., 2002) (1/4,000), and α-BAK1 (1/5,000; custom-made by Eurogentec). For the uncropped blots see Figure S3.

### Xylem staining

Seven-day-old Arabidopsis seedlings were stained with Basic Fuchsin as described (Ursache et al., 2018).

### Confocal microscopy and image analysis

For BRI1 localization Arabidopsis seedlings were imaged with an Olympus FluoView1000 confocal laser scanning microscope and a UPLSAPO 60×/1.2 n.a. water-corrected immersion objective at digital zoom 2, whereas for BRI1 internalization and BFA treatment, a Leica SP8X confocal microscope was used with a HC PL 584 APO CS2 40x/1.1 n.a. water-corrected immersion objective at digital zoom 5 and 3, respectively. The excitation/emission wavelengths were 514 nm/530-600 nm for BRI1-mCit. For the BRI1 internalization, the membrane of individual cells was selected using the brush tool of ImageJ with a size of 5 pixels as well as using the polygon selection tool to mark the intracellular space. The average intensity of the top 5 % highest pixels for both the PM and the intracellular space was used to obtain a ratio between PM and intracellular fluorescence. The BFA body size was calculated as previously described (Luo et al., 2015). Xylem was imaged with Leica SP8X confocal microscope equipped with a HC PL APO CS2 63x/1.20 n.a. water-corrected immersion objective. The excitation/emission wavelengths used were of 561 nm/600-650 nm for Basic Fuchsin staining. The BRI1 heat shock lines were analyzed under a vertical ZEISS LSM900 microscope equipped with a Plan-Apochromat M27 20x/ 0.8 n.a. objective. The excitation/emission wavelengths were 514 nm/530-600 nm for BRI1-mCit. The root tip was tracked over time with the TipTracker software (von Wangenheim et al., 2017). Quantification was obtained by measuring the PM signal of the same cell over time and normalized by the time point with the highest PM signal. Images processing and quantification were performed with the Fiji software package.

### Grating-coupled interferometry (GCI)

A Creoptix WAVE system (Creoptix AG, Switzerland) was used for the GCI binding assays. Experiments were done on a 4PCH WAVE GCI chip (long polycarboxylate surface; Creoptix AG). After a borate buffer conditioning (100 mM sodium borate, pH 9.0, 1 M NaCl; XanTec Bioanalytics, Düsseldorf, Germany), streptavidin was immobilized through a standard amine coupling protocol, followed by passivation of the surface (0.5% BSA [Roche] in 10 mM sodium acetate, pH 5.0), and final quenching with 1 M ethanolamine, pH 8.0 for 7 min (XanTec Bioanalytics). The LRR ectodomains of wild-type BRI1 and the respective mutants were biotinylated and coupled to the streptavidin-coated chip. For the BL binding experiments, BL was injected in a 1:2 dilution series, starting from 3 µM, in 20 mM citrate, pH 5.0, 250 mM NaCl at 25°C. Blank injections were used for double referencing and a DMSO calibration curve for bulk correction. All analyses and corrections were done with the Creoptix WAVE control software, with a one-to-one binding model with bulk correction used to fit all experiments.

### Analytical size-exclusion chromatography (SEC)

Analytical SEC experiments were carried out on a Superdex 200 increase 10/300 GL column (GE Healthcare), preequilibrated in 20 mM sodium citrate, pH 5.0, 250 mM NaCl. 200 µg of protein, injected in a 100-µl volume, was loaded onto the column and elution at 0.75 ml/min was monitored by ultraviolet absorbance at λ = 280 nm. Peak fractions were analyzed by SDS-PAGE.

### Right-angle light scattering (RALS)

BRI1 ectodomains (residues 1-788 with a C-terminal Avi-tag as well as a TEV protease cleavable TwinStrep – 9x His tag) were expressed and purified as described previously (Hohmann et al 2018b) and analyzed by size-exclusion chromatography (SEC) paired with a RALS and a refractive index (RI) detector, using an OMNISEC RESOLVE / REVEAL system. The calibration of the instrument was carried out with a BSA standard (Thermo Scientific). In a 50 µl volume, 100 µg of protein was separated on a Superdex 200 increase column (GE Healthcare) in 20 mM sodium citrate, pH 5.0, 250 mM NaCl at a column temperature of 35°C and a 0.7 ml/min. Data were analyzed using the OMNISEC software (v10.41).

### Homology modeling and structure visualization

BRI^Q^ ectodomain structure was modeled using the Modeller 9.18 program (Šali and Blundell, 1993) with the BRI1 ectodomain (PDB ID 3RJ0) as a template. The structures of wild type BRI1 and BRI1^Q^ ectodomains (Figure 1) were visualized by UCSF Chimera (Pettersen et al., 2021). Multiple sequence alignment was prepared using the Jalview program (Waterhouse et al., 2009)

### RT-qPCR

Seven-day-old seedlings in liquid half-strength Murashige and Skoog medium were transferred to 37°C for 1 h and let to recover at room temperature for another 1 h. Total RNA was extracted with the RNeasy kit (Qiagen). cDNA from RNA was synthesized with the qScript cDNA Supermix (Quantabio). RT-qPCRs were run with SYBR green I Master kit (Roche) on a LightCycler 480 (Roche). The mCitrine expression was normalized to that of ACTIN2 and GAPDH. The cycling conditions were as follows: preincubation at 95°C for 10 min; 45 amplification cycles at 95°C for 10 s, 60°C for 15 s, and 72°C for 15 s; melting curve at 95°C for 1 s, and 65°C for 1 s, followed by cooling at 40°C for 10 s.

## QUANTIFICATION AND STATISTICAL ANALYSIS

The data were subjected to statistical analysis using GraphPad Prism (https://www.graphpad.com/scientific-software/prism/) and Excel software. Comparisons between groups were made with the Mann Whitney test or two-way ANOVA with subsequent post hoc Sidak’s multiple comparisons test. Comparisons between discrete groups were made using Chi-square test. The measurements are shown as box plots displaying the first and third quartiles and split by medians (center lines), with whiskers extending to 1.5-fold the interquartile range from the 25th and 75th percentiles.

## SUPPLEMENTAL INFORMATION

Supplemental Information can be found online

## ACKNOWLEDGEMENTS

We thank Yanhai Yin (Iowa State University, Ames, USA), Gregory Vert (CNRS/Université de Toulouse, France), and Cyril Zipfel (University of Zurich, Switzerland) for providing the anti-BES1 antibody, the *pBRI1:BRI1-mCit*/*bri1 Arabidopsis* transgenic line, pDONRP4P1r-pBRI1, pDONRP1P2-BRI1 and pDONRP2rP3-mCit plasmids, and information for making the anti-BAK1 antibody, respectively, and Martine De Cock for help in preparing the manuscript. This work was supported by Ghent University Special Research Fund Grant (BOF15/24J/048 to E.R.), the Research Foundation-Flanders (project G022516N to E.R. and a postdoctroral fellowship 12R7819N to N.V.), the European Research Council (ERC Co T-Rex grant 682436 to D.V.D), the Swiss National Science Foundation (project number 31003A_176237, to M.H.), the Howard Hughes Medical Institute (to M.H.) and the German Research Foundation (DFG) (project WO 1660/6-2 to S.W.).

## AUTHOR CONTRIBUTIONS

L.A.N.C., D.L., N.V., and E.R. conceived, designed, and performed the research. R.P. did modeling. J. L., G.W., and Y.J. contributed unpublished materials. U.H., and M.H. did binding experiments. L.A.N.C., A.S. and S.W. did the xylem phenotype experiments. D.V.D. helped with data analysis and manuscript finalization. L.A.N.C., D.L., and E.R. wrote the manuscript. All authors commented on the results and on the manuscript text.

---

**Figure S1.**
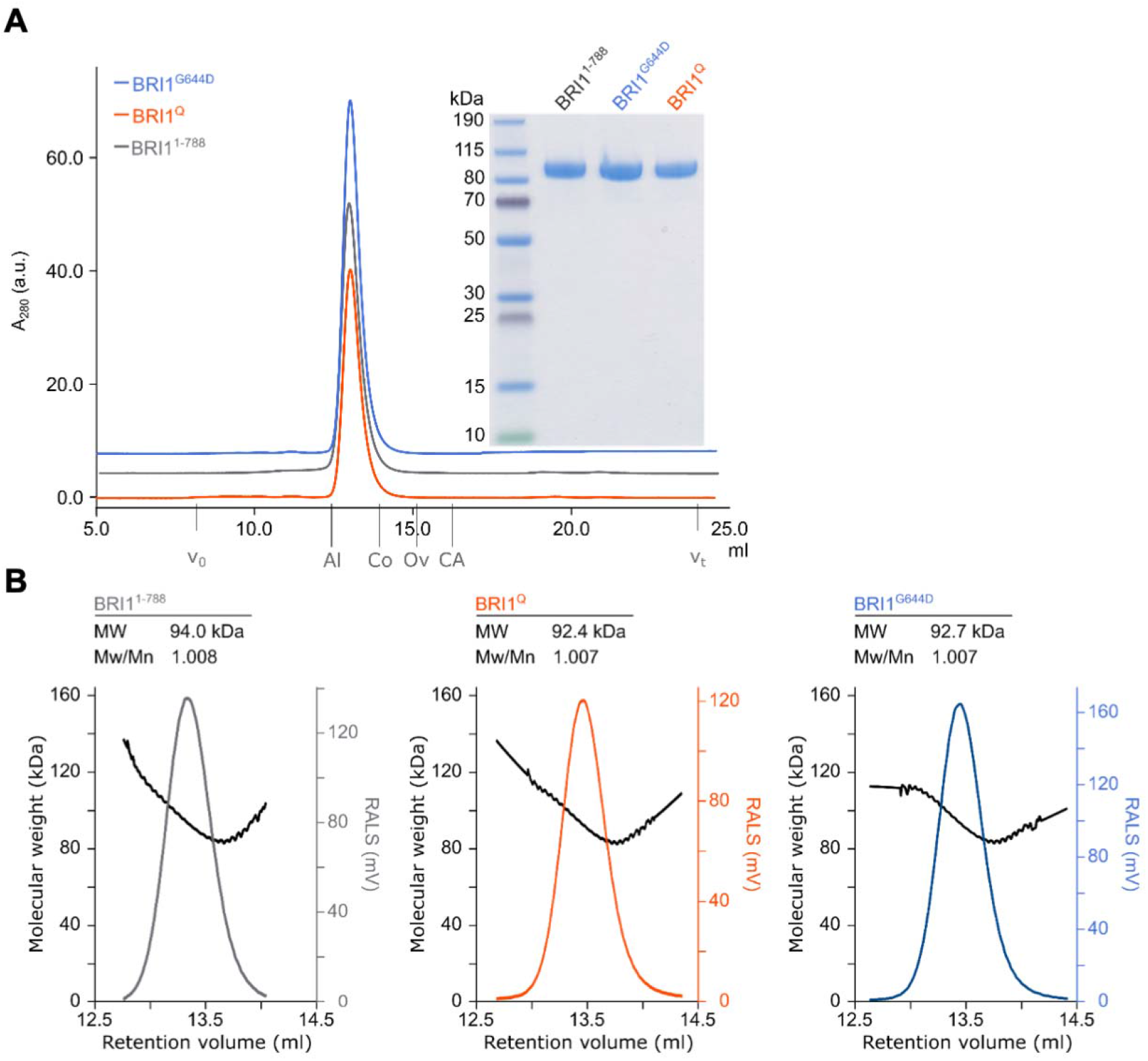
Purity and oligomeric state of wild-type and mutant BRI1 ectodomains. (A) Analytical size exclusion chromatography traces of BRI1^1-788^ (grey), BRI1^G644D^ (blue) and BRI1^Q^ (orange) with the SDS-PAGE analysis of pooled peak fractions alongside. Indicated are the void volume (v_0_), the total column volume (v_t_) and the elution volumes for molecular mass standards (aldolase [Al], 158 kDa; conalbumin [Co], 75 kDa; ovalbumin [Ov], 43 kDa; and carbonic anhydrase [Ca], 29 kDa). Coomassie-stained SDS-PAGE containing 5 μg of the indicated wild-type or mutated BRI1 ectodomains were isolated from monomeric peak fractions purified by size-exclusion chromatography. (B) Analysis of the oligomeric state of BRI1 ectodomains. Shown are raw right-angle light scattering traces (grey, blue, and orange) and extrapolated molecular weight (black) of BRI1^1-788^, BRI1^Q^, and BRI1^G644D^, including observed molecular weight (MW) and the dispersity (Mw/Mn). The theoretical molecular weight is 83.5 kDa for BRI1^1-788^, 83.3 kDa for BRI1^Q^ and 83.6 kDa for BRI1^G644D^.

**Figure S2.**
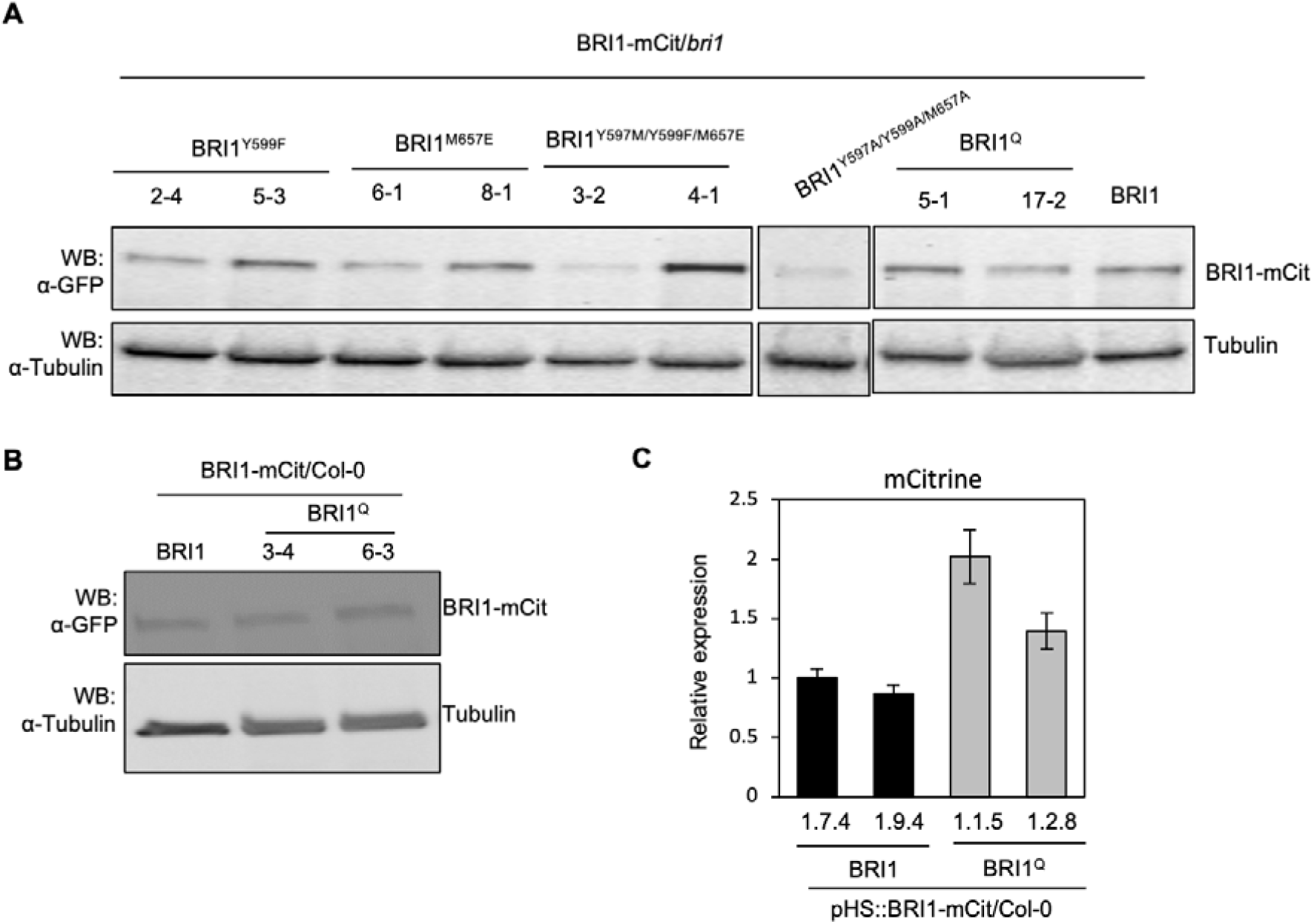
BRI1 expression in different BRI1 transgenic lines. (A and B) Total proteins isolated from 5-day-old seedlings and detected by Western blot (WB) with α-GFP antibodies to detect the BRI1-mCitrine (mCit). Numbers indicate individual lines. α-tubulin antibodies were used to show equilibrate protein loading. (C) Real-time quantitative reverse transcription PCR (qRT-PCR) analysis of mCitrine. Total RNA was isolated from 7-day-old seedlings of two independent transgenic lines. The seedlings in liquid half-strength Murashige and Skoog medium were transferred to 37°C for h and left to recover at room temperature for 1 h. *GAPDH* and *ACTIN2* were used as normalization controls. In the graph, the mean ± SD (*n*=3) is presented.

**Figure S3.**
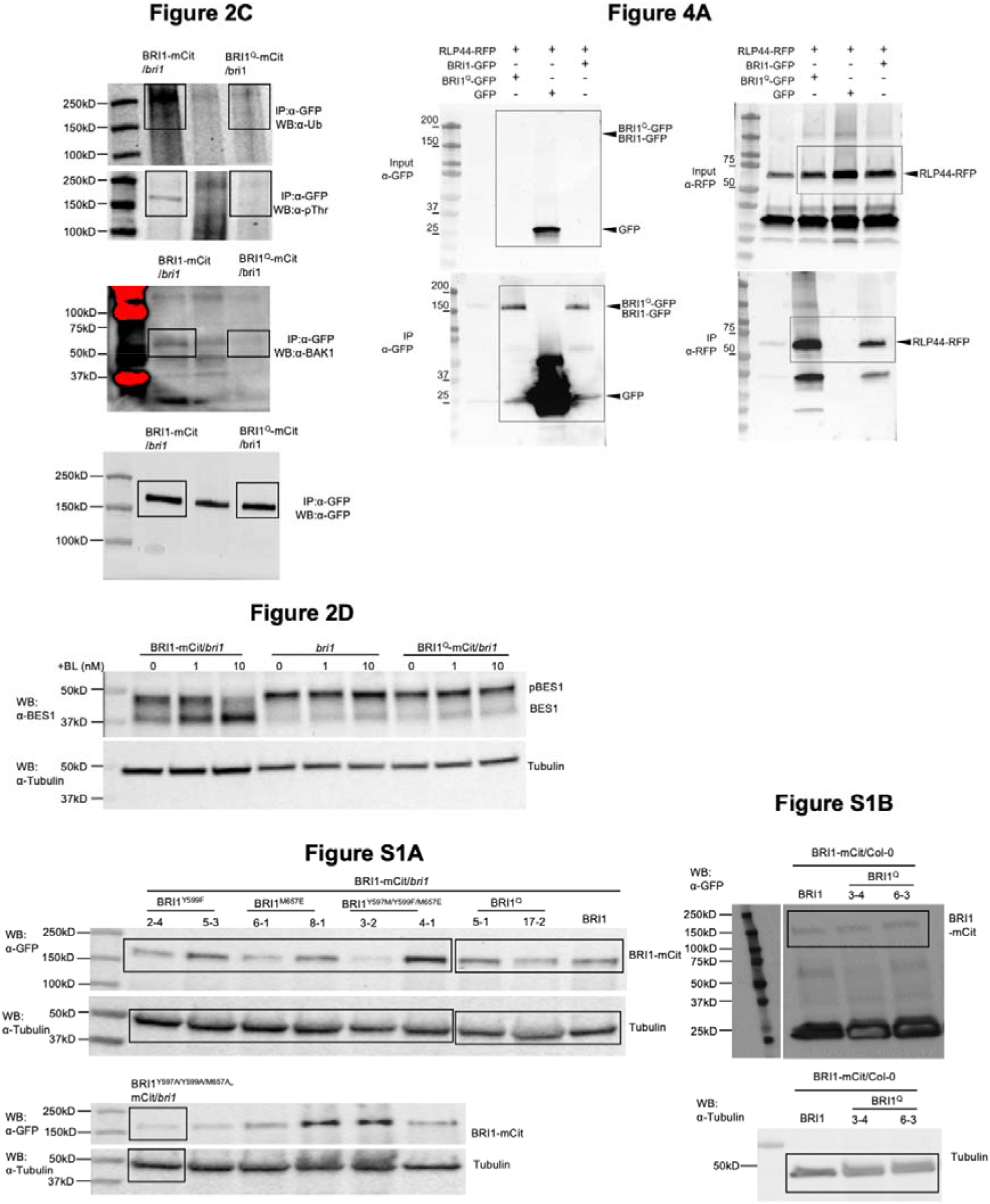
Source blots.

**Table S1.**
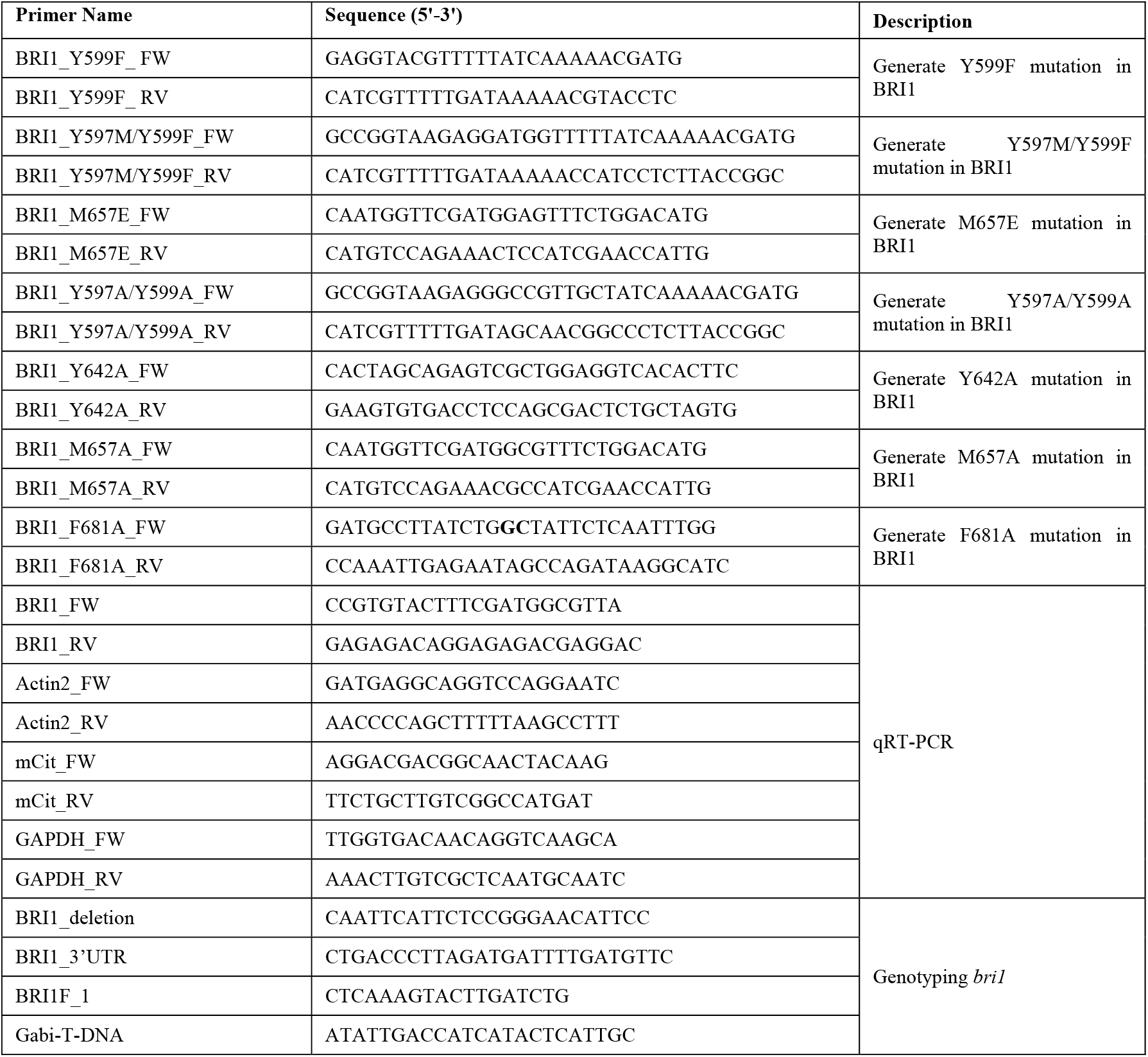
List of oligonucleotides used in this study.

